# Modeling Single-Cell Dynamics Using Unbalanced Parameterized Monge Maps

**DOI:** 10.1101/2022.10.04.510766

**Authors:** Luca Vincent Eyring, Dominik Klein, Giovanni Palla, Soeren Becker, Philipp Weiler, Niki Kilbertus, Fabian J. Theis

**Author notes:** equal contribution (Luca Vincent Eyring, Dominik Klein, Giovanni Palla).

## Abstract

Optimal Transport (OT) has proven useful to infer single-cell trajectories of developing biological systems by aligning distributions across time points. Recently, Parameterized Monge Maps (PMM) were introduced to learn the optimal map between two distributions. Here, we apply PMM to model single-cell dynamics and show that PMM fails to account for asymmetric shifts in cell state distributions. To alleviate this limitation, we propose Unbalanced Parameterised Monge Maps (UPMM). We first describe the novel formulation and show on synthetic data how our method extends discrete unbalanced OT to the continuous domain. Then, we demonstrate that UPMM outperforms well-established trajectory inference methods on real-world developmental single-cell data.

## 1 Introduction

Single-cell RNA sequencing (scRNA-seq) data allows to study cellular development at an unprecedented resolution. To recover these developmental landscapes, various trajectory inference (TI) methods have been proposed [Sae+19]. RNA velocity-based methods ([La +18], [Ber+20]) derive the developmental dynamics by modeling spliced and unspliced gene expression rates with ordinary differential equations, yielding velocity vectors in gene space for every single cell. Optimal Transport (OT, [PC+19]) has also proven useful for inferring developmental dynamics in temporal scRNA-seq data[Sch+19]. However, this approach is limited by its computational complexity and only yields couplings between cell populations as opposed to generating vectors in gene expression space. In this work, we alleviate these limitations by suggesting a TI algorithm based on Optimal Transport via Input Convex Neural Networks (ICNNs, [AXK17]), which we refer to as Parameterized Monge Maps (PMM) [Mak+20]). PMM has been successfully applied to modeling perturbation responses in scRNA-seq data [Bun+21; BKC22]. However, one major disadvantage of applying PMM to scRNA-seq data is the lack of an unbalanced formulation, allowing the model to adapt the source and target distribution during training. This concept is needed in many applications, e.g. to account for different proliferation rates of cells.

We adapt PMM in two different ways to overcome this limitation. First, we show how to adapt the distribution based on prior biological knowledge to allow for different proliferation rates, which has been successfully used in the discrete OT case [Sch+19]. Second, we propose Unbalanced Parameterized Monge Maps, a novel algorithm based on PMM. On simulated data, we demonstrate that it mimics the behavior of discrete unbalanced OT. Finally, we apply our algorithms to a scRNAseq dataset of pancreatic endocrinogenesis [Bas+19] and disentangle fine-grained cell state trajectories. We show that our proposed methods outperform the established trajectory inference methods scVelo [Ber+20], Waddington OT [Sch+19], and TrajectoryNet [Ton+20].

## 2 Methods

Let *P* and *Q* be two probability distributions defined on ℝ^*d*^. Let *X* and *Y* be random variables such that *X* ~ *P* and *Y* ~ *Q*, respectively. The Monge Map is defined as

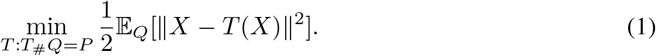

If distances in the space are measured in the squared Euclidean distance, this optimization problem can be rewritten to ([Vil03], Theorem 1.3)

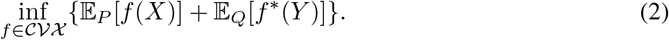

where *𝒞𝒱𝒳* is the set of integrable convex functions and *f* ^*\**^ denotes the convex conjugate of *f* defined by *f \**(*y*)= sup_*x*_⟨*x, y*⟩− *f*(*x*). Moreover, the Monge Map *T* : *Q* → *P* can be obtained as the gradient of the convex function *f* ^*^:

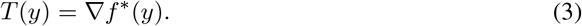

Thus, Makkuva et al. propose to learn *f* and *g* with Input Convex Neural Networks (ICNNs, [AXK17]) leading to the optimization problem

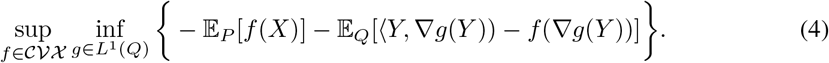

Thus, we obtain a fully parameterized vector field by taking the gradient of the potential *g* since *T* (*y*)= ∇ *g*(*y*). See Appendix A.1 for a detailed derivation.

### 2.1 PMM with transformed distributions

For modeling scRNA-seq data capturing different developmental stages, a notion of unbalancedness is crucial as some cells might proliferate faster than others. Schiebinger et al. propose using proliferation and apoptosis genes to model cell growth and death, respectively, in the discrete OT case. Similarly, we transform the source distribution *Q* with a function γ : *Q* → *Q* depending on proliferation and apoptosis marker genes. Hence, we solve (1) with *X* ~ γ(*Q*) (Appendix A.12).

If prior knowledge is not available, γ cannot be defined. Consequently, PMM cannot account for unbalanced distribution shifts (e.g. across time points). To alleviate this limitation, we propose Unbalanced Parameterized Monge Maps (UPMM).

### 2.2 Unbalanced Parameterized Monge Maps

The goal of UPMM is to mimic the behavior of discrete unbalanced optimal transport (Appendix A.2) in the continuous domain. Therefore, we define the objective function of UPMM as

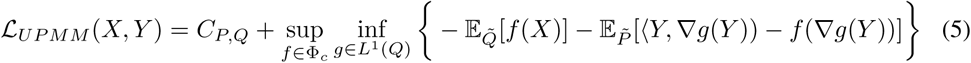

Here, 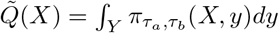 (*X, y*)*dy* and 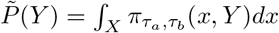, where 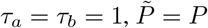denotes the Wasserstein-2 optimal coupling of *X* and *Y*. We estimate 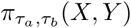 batch-wise with discrete regularised OT [Cut13]. Analogous to the discrete unbalanced OT formulation, decreasing τ_*a*_ and τ_*b*_ increases unbalancedness in source and target distribution, respectively^3^. If 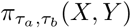 and 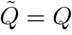, hence we recover PMM as defined in (4). Algorithm A.3 shows the full training procedure, with training details and hyperparameters described in Appendix A.4.

For simulated data (Appendix A.5.1), Figure 1 visually confirms that UPMM with τ = τ_*a*_ = τ_*b*_ =1 yields the same results as PMM. By gradually decreasing τ, we arrive at a map with similar behavior as the one obtained from the discrete unbalanced case. The corresponding potentials are shown in Appendix A.11.1.

**Figure 1:**
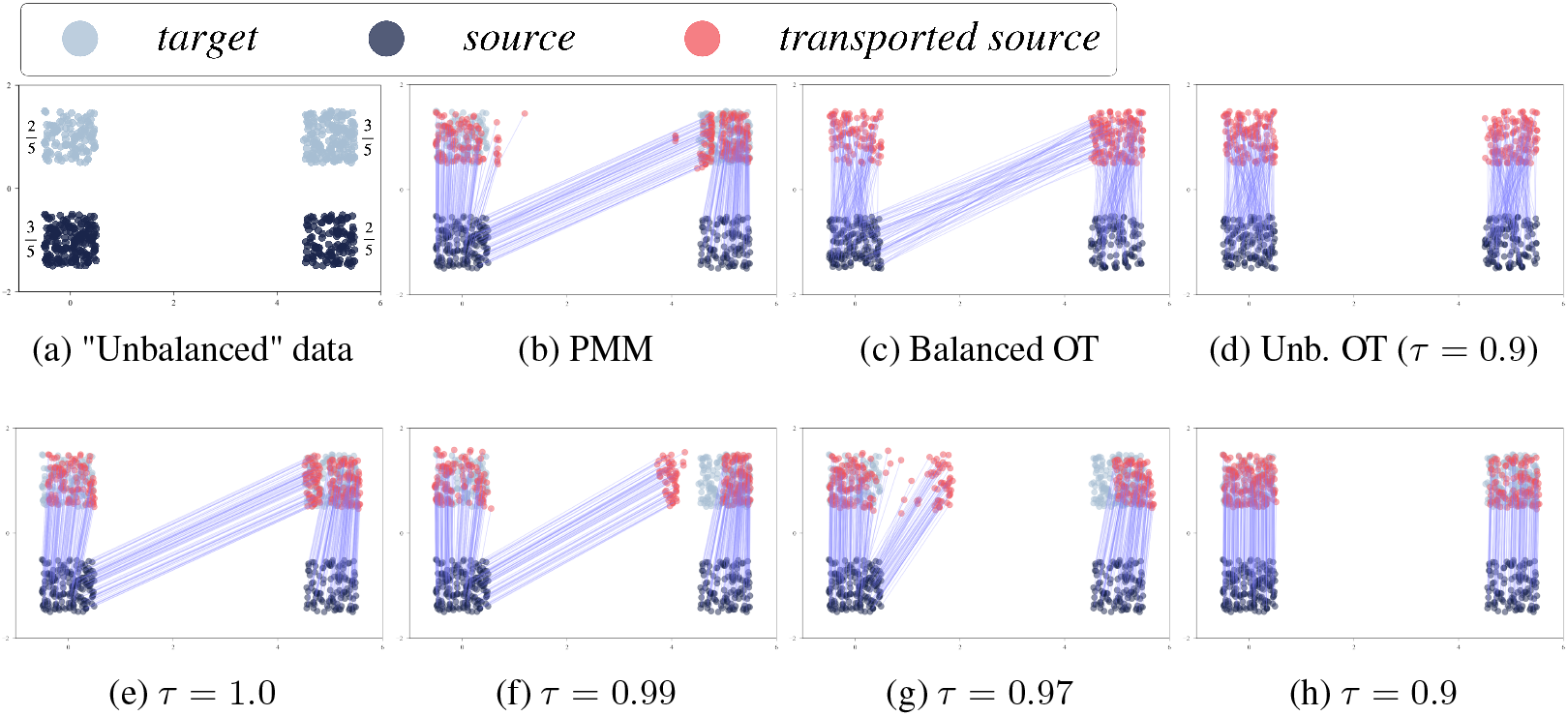
Different maps on data drawn from a mixture of uniform distribution, where the density in the bottom left 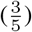 and the top right 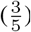 is higher then in the top left 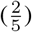 and bottom right 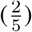, (Appendix A.5.1). Besides the data in Figure 1a, the first row shows results of PMM as proposed in [Mak+20](1b), discrete balanced OT (1c), and discrete unbalanced OT (1d). The second row shows the maps obtained by UPMM with different unbalancedness parameters τ.

### 2.3 Benchmarks and metrics

To assess the performance of PMM, UPMM, and PMM with growth rate prior (Appendix A.12), which we refer to as PMM+GR, we apply our three newly proposed TI methods to a pancreatic endocrinogenesis scRNA-seq dataset (Appendix A.5.2). We compare our methods to the wellestablished TI methods scVelo, Waddington OT, and TrajectoryNet (Appendix A.6). To evaluate how accurately different methods recover biological ground truth, we rely on CellRank [Lan+22] to quantify cell fate probabilities.

The performance of all methods is assessed with three metrics. First, we consider aggregated *cell type transitions* based on biological knowledge. We consider cell type transitions in three classes, the endocrine branch (EB), Ngn3 EP, and non-endocrine branch (NEB) (Appendix A.8.1), which we refer to as lineages in the following. Here, we report the mean of correct cell transitions for each lineage, while full results can be found in Appendix A.8. Second, *cell type redundancy* determines to which extent terminal cell populations are recovered with CellRank (Appendix A.9). This quantifies how well terminal cell states can be recovered from the learned dynamics. Third, we analyze *velocity consistency* across transcriptomically similar cells (Appendix A.10) justified by the fact that cells that are close in gene expression space should have a similar cell trajectory.

## 3 Results

Table 1 shows that, overall, our newly proposed TI methods PMM+GR and UPMM recover correct cell type transitions best. None of the competing methods yield comparable results across all considered sets of cell types. TrajectoryNet maps most Ngn3 high endocrine progenitors (Ngn3 EP) incorrectly. Appendix A.13 reveals that most of these cells are mapped to Acinar cells. While scVelo performs well on Ngn3 EP cells it fails on various other cell types, for example having a large bias to map EP cells to Beta cells. WOT accurately recovers the NEB lineage but fails to achieve similar performance on the other two lineages. Table 1 shows that PMM yields promising results in the EB lineage but its performance on Ngn3 EP is very poor. This can be explained by the asymmetric distribution shift of cell states between the two time points. In effect, Acinar cells proliferate much faster than cells in the EB lineage as can be seen in Appendix A.5.2. Hence, Ngn3 EP is mainly mapped to the NEB branch. In contrast, PMM+GR can compensate for that by considering proliferation and apoptosis genes. Hence, almost all Ngn3 (EP) cells are mapped correctly. Similarly, UPMM explicitly models unbalancedness, thus ranking second overall. While table 1 reports aggregated results, table 2 in Appendix A.8.2 shows more detailed results.

**Table 1:**
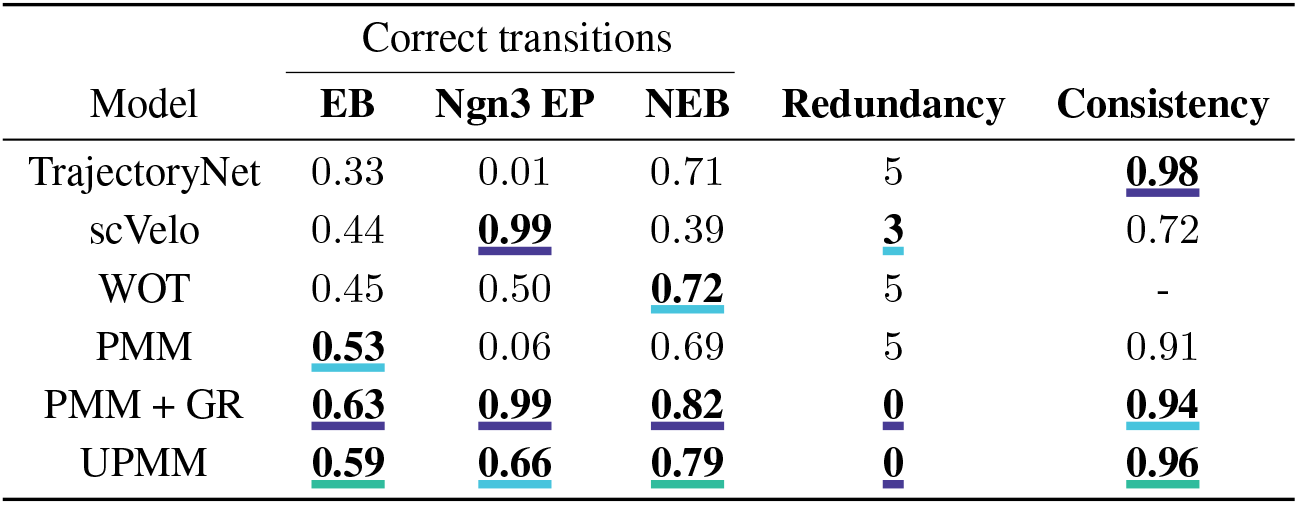
Evaluation of different trajectory inference methods based on three different metrics. Columns 1-3 show the fraction of correct cell type transitions, column 4 contains the cell type redundancy score, and column 5 shows the mean velocity consistency. For each column, we underline the best, second best, and third best methods.

Figure 2 complements these quantitative results by projecting the high dimensional velocities onto a two-dimensional UMAP [MHM18] representation (Appendix A.7). For example, the velocity flow estimated by PMM contradicts the biological ground truth for the Ngn3 EP cells, whose velocity vectors should point towards the upper left of the UMAP in (2c). While UPMM accounts considerably well for the unbalancedness (2d), PMM+GR yields optimal directionalities of Ngn3 EP cells (2e). Plots of the potential values of the UMAP-embedded cells can be found in Appendix A.11.2.

**Figure 2:**
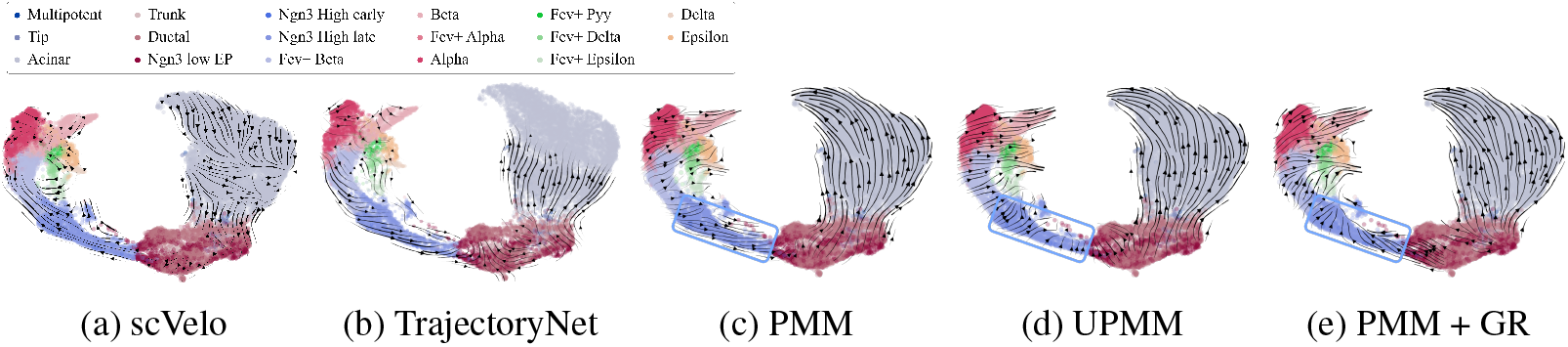
Velocity stream embedding plots. For our newly proposed TI methods (Figures 2c, 2d, 2e) the blue box highlight the direction of Ngn3 EP cells, demonstrating the need of incorporating unbalancedness.

The superior performance of PMM+GR and UPMM is confirmed by the cell type redundancy metric in Table 1. They both yield optimal matches between recovered and known terminal states. Table 1 also shows that velocity vectors of similar cells are significantly higher correlated for PMM, UPMM, PMM+GR, and TrajectoryNet than for scVelo. WOT does not yield velocity vectors and hence neither can the velocity consistency metric be computed nor the velocity stream embedding be plotted (Appendix A.10). Overall, our proposed methods outperform well-established trajectory inference methods. While PMM is a promising approach to model cell dynamics we showed that incorporating unbalancedness is crucial for the accurate identification of cell fate maps.

## 4 Discussion

By extending PMM to the unbalanced setting we propose a new, powerful trajectory inference algorithm that is able to recover fine-grained cell fates while being robust to unbalanced population growth of cells. Given its ability to learn a generative continuous map of cellular differentiation, we believe PMM+GR and UPMM would also show competitive performance for interpolating cellular states to unseen cell types during training. Our methods are particularly valuable for development cell atlas building ([Han+21]) due to their performance, scalability and applicability to unseen data during training. Moreover, our algorithm naturally extends to multiple modalities, e.g. allowing for a fully parameterized vector field in chromatin or protein space. Instead of seeing RNA velocity and UPMM as competing approaches, a promising approach would be to superpose the velocity vector fields to combine strengths from both methods.

## Footnotes

3 We follow the notation in [Cut+22b], where 0 < τ*i* ≤1 with τ_*i*_ = 1 corresponding to the balanced case (Appendix A.2).

## Notes

### Competing Interest Statement

Fabian J. Theis consults for Immunai Inc., Singularity Bio B.V., CytoReason Ltd, and Omniscope Ltd, and has ownership interest in Dermagnostix GmbH and Cellarity.

## References

[Vil03] C. Villani. Topics in Optimal Transportation. Graduate studies in mathematics. American Mathematical Society, 2003. ISBN: 9780821833124. URL: https://books.google.es/books?id=R%5C_nWqjq89oEC.

[Cut13] Marco Cuturi. “Sinkhorn Distances: Lightspeed Computation of Optimal Transport”. In: Advances in Neural Information Processing Systems. Ed. by C.J. Burges et al. Vol. 26. Curran Associates, Inc., 2013. URL: https://proceedings.neurips.cc/paper/2013/file/af21d0c97db2e27e13572cbf59eb343d-Paper.pdf.

[Tir+16] Itay Tirosh et al. “Single-cell RNA-seq supports a developmental hierarchy in human oligodendroglioma”. In: Nature 539.7628 (2016), pp. 309–313.

[AXK17] Brandon Amos, Lei Xu, and J Zico Kolter. “Input convex neural networks”. In: International Conference on Machine Learning. PMLR. 2017, pp. 146–155.

[Bra+18] James Bradbury et al. JAX: composable transformations of Python+NumPy programs. Version 0.3.13. 2018. URL: http://github.com/google/jax.

[Chi+18] Lenaic Chizat et al. “Unbalanced optimal transport: Dynamic and Kantorovich formulations”. In: Journal of Functional Analysis 274.11 (2018), pp. 3090–3123.

[La +18] Gioele La Manno et al. “RNA velocity of single cells”. en. In: Nature 560.7719 (Aug. 2018), pp. 494–498.

[MHM18] Leland McInnes, John Healy, and James Melville. UMAP: Uniform Manifold Approximation and Projection for Dimension Reduction. 2018. DOI: 10.48550/ARXIV.1802.03426. URL: https://arxiv.org/abs/1802.03426.

[WAT18] F Alexander Wolf, Philipp Angerer, and Fabian J Theis. “SCANPY: large-scale single-cell gene expression data analysis”. In: Genome biology 19.1 (2018), pp. 1–5.

[Bas+19] Aimée Bastidas-Ponce et al. “Comprehensive single cell mRNA profiling reveals a detailed roadmap for pancreatic endocrinogenesis”. In: Development 146.12 (2019), dev173849.

[Kor+19] Alexander Korotin et al. “Wasserstein-2 Generative Networks”. In: CoRR abs/1909.13082 (2019). arXiv: 1909.13082. URL: http://arxiv.org/abs/1909.13082.

[PC+19] Gabriel Peyré, Marco Cuturi, et al. “Computational optimal transport: With applications to data science”. In: Foundations and Trends® in Machine Learning 11.5-6 (2019), pp. 355–607.

[Sae+19] Wouter Saelens et al. “A comparison of single-cell trajectory inference methods”. In: Nature biotechnology 37.5 (2019), pp. 547–554.

[Sch+19] Geoffrey Schiebinger et al. “Optimal-transport analysis of single-cell gene expression identifies developmental trajectories in reprogramming”. In: Cell 176.4 (2019), pp. 928– 943.

[Bab+20] Igor Babuschkin et al. The DeepMind JAX Ecosystem. 2020. URL: http://github.com/deepmind.

[Ber+20] Volker Bergen et al. “Generalizing RNA velocity to transient cell states through dynamical modeling”. en. In: Nat. Biotechnol. (Aug. 2020).

[Mak+20] Ashok Makkuva et al. “Optimal transport mapping via input convex neural networks”. In: International Conference on Machine Learning. PMLR. 2020, pp. 6672–6681.

[Ton+20] Alexander Tong et al. “Trajectorynet: A dynamic optimal transport network for modeling cellular dynamics”. In: International conference on machine learning. PMLR. 2020, pp. 9526–9536.

[Bun+21] Charlotte Bunne et al. “Learning Single-Cell Perturbation Responses using Neural Optimal Transport”. In: bioRxiv (2021).

[Han+21] Muzlifah Haniffa et al. “A roadmap for the Human Developmental Cell Atlas”. en. In: Nature 597.7875 (Sept. 2021), pp. 196–205.

[Amo+22] Brandon Amos et al. Meta Optimal Transport. 2022. DOI: 10.48550/ARXIV.2206.05262. URL: https://arxiv.org/abs/2206.05262.

[BKC22] Charlotte Bunne, Andreas Krause, and Marco Cuturi. “Supervised Training of Conditional Monge Maps”. In: arXiv preprint arXiv:2206.14262 (2022).

[Cut+22a] Marco Cuturi et al. Optimal Transport Tools (OTT): A JAX Toolbox for all things Wasserstein. 2022. DOI: 10.48550/ARXIV.2201.12324. URL: https://arxiv.org/abs/2201.12324.

[Cut+22b] Marco Cuturi et al. “Optimal transport tools (ott): A jax toolbox for all things wasserstein”. In: arXiv preprint arXiv:2201.12324 (2022).

[Lan+22] Marius Lange et al. “CellRank for directed single-cell fate mapping”. en. In: Nat. Methods 19.2 (Feb. 2022), pp. 159–170.

