## Appendix for "Modeling Single-Cell Dynamics Using Unbalanced Parameterized Monge Maps"


**Luca Vincent Eyring***
Institute of Computational Biology
Helmholtz Center Munich
Germany

**Dominik Klein***
Institute of Computational Biology
Helmholtz Center Munich
Germany

**Giovanni Palla***
Institute of Computational Biology
Helmholtz Center Munich
School of Life Sciences Weihenstephan
Technical University of Munich Germany

**Soeren Becker**
Helmholtz AI, Munich
Germany

**Philipp Weiler**
Institute of Computational Biology
Helmholtz Center Munich
Department of Mathematics
Technical University of Munich Germany

**Niki Kilbertus**
Helmholtz AI, Munich
Technical University of Munich
Germany

**Fabian J. Theis** †
Institute of Computational Biology
Helmholtz Center Munich
School of Life Sciences Weihenstephan
Department of Mathematics
Technical University of Munich Germany



#### Abstract

Optimal Transport (OT) has proven useful to infer single-cell trajectories of developing biological systems by aligning distributions across time points. Recently, Parameterized Monge Maps (PMM) were introduced to learn the optimal map between two distributions. Here, we apply PMM to model single-cell dynamics and show that PMM fails to account for asymmetric shifts in cell state distributions. To alleviate this limitation, we propose Unbalanced Parameterised Monge Maps (UPMM). We first describe the novel formulation and show on synthetic data how our method extends discrete unbalanced OT to the continuous domain. Then, we demonstrate that UPMM outperforms well-established trajectory inference methods on real-world developmental single-cell data.


#### 1 Introduction

Single-cell RNA sequencing (scRNA-seq) data allows to study cellular development at an unprecedented resolution. To recover these developmental landscapes, various trajectory inference (TI)


*equal contribution
†corresponding:

#### 2 Methods

Let $P$ and $Q$ be two probability distributions defined on $\mathbb{R}^d$. Let $X$ and $Y$ be random variables such that $X \sim P$ and $Y \sim Q$, respectively. The Monge Map is defined as

$$\min_{T:T_{\#}Q=P} \frac{1}{2}\mathbb{E}_Q[\|X - T(X)\|^2]. \tag{1}$$

If distances in the space are measured in the squared Euclidean distance, this optimization problem can be rewritten to ([Vil03], Theorem 1.3)

$$\inf_{f \in \mathcal{CVX}} \{\mathbb{E}_P[f(X)] + \mathbb{E}_Q[f^*(Y)]\}. \tag{2}$$

where $\mathcal{CVX}$ is the set of integrable convex functions and $f^*$ denotes the convex conjugate of $f$ defined by $f^*(y) = \sup_x \langle x, y \rangle - f(x)$. Moreover, the Monge Map $T : Q \to P$ can be obtained as the gradient of the convex function $f^*$:

$$T(y) = \nabla f^*(y). \tag{3}$$

Thus, Makkuva et al. propose to learn $f$ and $g$ with Input Convex Neural Networks (ICNNs, [AXK17]) leading to the optimization problem

$$\sup_{f \in \mathcal{CVX}} \inf_{g \in L^1(Q)} \left\{ -\mathbb{E}_P[f(X)] - \mathbb{E}_Q[\langle Y, \nabla g(Y)) - f(\nabla g(Y))] \right\}. \tag{4}$$

Thus, we obtain a fully parameterized vector field by taking the gradient of the potential $g$ since $T(y) = \nabla g(y)$. See Appendix A.1 for a detailed derivation.

##### 2.1 PMM with transformed distributions

For modeling scRNA-seq data capturing different developmental stages, a notion of unbalancedness is crucial as some cells might proliferate faster than others. Schiebinger et al. propose using proliferation and apoptosis genes to model cell growth and death, respectively, in the discrete OT case. Similarly, we transform the source distribution $Q$ with a function $\gamma : Q \to Q$ depending on proliferation and apoptosis marker genes. Hence, we solve (1) with $X \sim \gamma(Q)$ (Appendix A.12).

If prior knowledge is not available, $\gamma$ cannot be defined. Consequently, PMM cannot account for unbalanced distribution shifts (e.g. across time points). To alleviate this limitation, we propose Unbalanced Parameterized Monge Maps (UPMM).

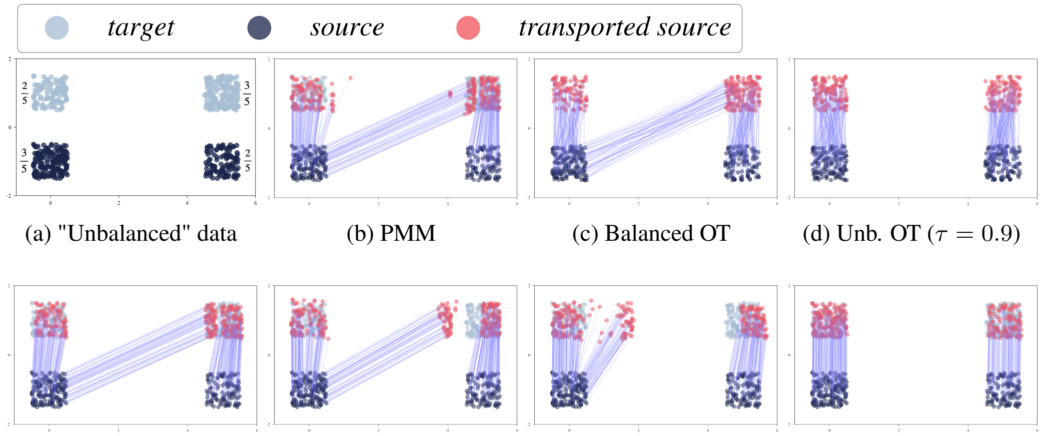


(a) "Unbalanced" data (b) PMM (c) Balanced OT (d) Unb. OT ($\tau = 0.9$)

(e) $\tau = 1.0$ (f) $\tau = 0.99$ (g) $\tau = 0.97$ (h) $\tau = 0.9$

Figure 1: Different maps on data drawn from a mixture of uniform distribution, where the density in the bottom left ($\frac{3}{5}$) and the top right ($\frac{3}{5}$) is higher then in the top left ($\frac{2}{5}$) and bottom right ($\frac{2}{5}$), (Appendix A.5.1). Besides the data in Figure 1a, the first row shows results of PMM as proposed in [Mak+20](1b), discrete balanced OT (1c), and discrete unbalanced OT (1d). The second row shows the maps obtained by UPMM with different unbalancedness parameters $\tau$.

##### 2.2 Unbalanced Parameterized Monge Maps

The goal of UPMM is to mimic the behavior of discrete unbalanced optimal transport (Appendix A.2) in the continuous domain. Therefore, we define the objective function of UPMM as

$$\mathcal{L}_{UPMM}(X,Y) = C_{P,Q} + \sup_{f\in\Phi_c} \inf_{g\in L^1(Q)} \Big\{ - \mathbb{E}_{\tilde{Q}}[f(X)] - \mathbb{E}_{\tilde{P}}[\langle Y, \nabla g(Y)\rangle - f(\nabla g(Y))] \Big\} \quad (5)$$

Here, $\tilde{Q}(X) = \int_Y \pi_{\tau_a,\tau_b}(X,y)dy$ and $\tilde{P}(Y) = \int_X \pi_{\tau_a,\tau_b}(x,Y)dx$, where $\pi_{\tau_a,\tau_b}(X,Y)$ denotes the Wasserstein-2 optimal coupling of $X$ and $Y$. We estimate $\pi_{\tau_a,\tau_b}(X,Y)$ batch-wise with discrete regularised OT [Cut13]. Analogous to the discrete unbalanced OT formulation, decreasing $\tau_a$ and $\tau_b$ increases unbalancedness in source and target distribution, respectively[3]. If $\tau_a = \tau_b = 1$, $\tilde{P} = P$ and $\tilde{Q} = Q$, hence we recover PMM as defined in (4). Algorithm A.3 shows the full training procedure, with training details and hyperparameters described in Appendix A.4.

For simulated data (Appendix A.5.1), Figure 1 visually confirms that UPMM with $\tau = \tau_a = \tau_b = 1$ yields the same results as PMM. By gradually decreasing $\tau$, we arrive at a map with similar behavior as the one obtained from the discrete unbalanced case. The corresponding potentials are shown in Appendix A.11.1.

##### 2.3 Benchmarks and metrics

To assess the performance of PMM, UPMM, and PMM with growth rate prior (Appendix A.12), which we refer to as PMM+GR, we apply our three newly proposed TI methods to a pancreatic endocrinogenesis scRNA-seq dataset (Appendix A.5.2). We compare our methods to the well-established TI methods scVelo, Waddington OT, and TrajectoryNet (Appendix A.6). To evaluate how accurately different methods recover biological ground truth, we rely on CellRank [Lan+22] to quantify cell fate probabilities.

The performance of all methods is assessed with three metrics. First, we consider aggregated *cell type transitions* based on biological knowledge. We consider cell type transitions in three classes, the endocrine branch (EB), Ngn3 EP, and non-endocrine branch (NEB) (Appendix A.8.1), which we refer to as lineages in the following. Here, we report the mean of correct cell transitions for each lineage, while full results can be found in Appendix A.8. Second, *cell type redundancy* determines to

[3] We follow the notation in [Cut+22b], where $0 < \tau_i \leq 1$ with $\tau_i = 1$ corresponding to the balanced case (Appendix A.2).

Table 1: Evaluation of different trajectory inference methods based on three different metrics. Columns 1-3 show the fraction of correct cell type transitions, column 4 contains the cell type redundancy score, and column 5 shows the mean velocity consistency. For each column, we underline the best, second best, and third best methods.

| | Correct transitions | | | | |
|---|---|---|---|---|---|
| Model | **EB** | **Ngn3 EP** | **NEB** | **Redundancy** | **Consistency** |
| TrajectoryNet | 0.33 | 0.01 | 0.71 | 5 | **0.98** |
| scVelo | 0.44 | **0.99** | 0.39 | **3** | 0.72 |
| WOT | 0.45 | 0.50 | **0.72** | 5 | - |
| PMM | **0.53** | 0.06 | 0.69 | 5 | 0.91 |
| PMM + GR | **0.63** | **0.99** | **0.82** | **0** | **0.94** |
| UPMM | **0.59** | **0.66** | **0.79** | **0** | **0.96** |

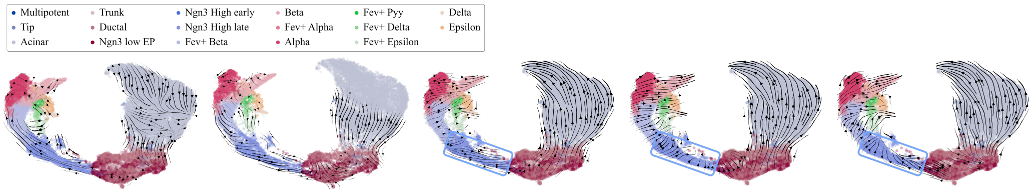


(a) scVelo (b) TrajectoryNet (c) PMM (d) UPMM (e) PMM + GR

Figure 2: Velocity stream embedding plots. For our newly proposed TI methods (Figures 2c, 2d, 2e) the blue box highlight the direction of Ngn3 EP cells, demonstrating the need of incorporating unbalancedness.

[Lan+22] Marius Lange et al. "CellRank for directed single-cell fate mapping". en. In: *Nat. Methods* 19.2 (Feb. 2022), pp. 159–170.

### A Appendix

#### A.1 Background on Monge Maps

In the following, we provide more details on the derivation of the objective function (4). The existence of the solution of the Monge Map (1) is not guaranteed [Mak+20], hence the Kantorovich relaxation is usually employed, defined as

$$W_2^2(P,Q) = \inf_{\pi \in \Pi(X,Y)} \int \|x-y\|_2^2 d\pi(x,y), \tag{6}$$

where $\Pi(X,Y)$ is the set of all joint probability distributions of $P$ and $Q$. In the discrete case, this problem is often solved with a regularized approach ([Cut13]) that is computationally more efficient.

The OT problem defined in (6) allows for a dual formulation [PC+19]

$$W_2^2(X,Y) = \sup_{f,g \in \Phi_c} \mathbb{E}_P[f(X)] + \mathbb{E}_Q[g(Y)], \tag{7}$$

where $\Phi_c = \{(f,g) \in L^1(P) \times L^1(Q) : f(x) + g(y) \leq \frac{1}{2}\|x-y\|_2^2 \,\forall (x,y) \in dP \otimes dQ \text{ a.e.}\}$. Villani ([Vil03], Theorem 1.3) reformulates (7) to

$$W_2^2(X,Y) = \frac{1}{2}\mathbb{E}_{P\otimes Q}[\|X\|_2^2 + \|Y\|_2^2] - \inf_{f \in \tilde{\Phi}_c} \{\mathbb{E}_P[f(X)] + \mathbb{E}_Q[f^*(Y)]\}. \tag{8}$$

where $\tilde{\Phi}_c = \{(f,g) \in L^1(P) \times L^1(Q) : f(x) + g(y) \geq \frac{1}{2}\langle f,g\rangle \,\forall (x,y) \in dP \otimes dQ \text{ a.e.}\}$. $f^*$ denotes the convex conjugate of $f$ defined by $f^*(y) = \sup_x \langle x,y\rangle - f(x)$. Note that $C_{P,Q} := \mathbb{E}_{P\otimes Q}[\|X\|_2^2 + \|Y\|_2^2]$ is constant in $f$. Recent advances propose to learn the optimal transport map in a fully parametric way by leveraging neural networks[Mak+20; Kor+19]. Makkuva et al. ([Mak+20]) make use of the fact that $\langle y, \nabla g(y)\rangle - f(\nabla g(y)) \leq f^*(y)$ for all functions $g : \mathbb{R}^d \to \mathbb{R}$ with equality attainable (with $g = f^*$) such that

$$W_2^2(X,Y) = C_{P,Q} + \sup_{f \in \tilde{\Phi}_c} \inf_{g \in L^1(Q)} \Big\{ -\mathbb{E}_P[f(X)] - \mathbb{E}_Q[\langle Y, \nabla g(Y)) - f(\nabla g(Y))] \Big\}. \tag{9}$$

#### A.2 Unbalanced discrete Optimal Transport

In this section, we introduce the notation for the balanced and unbalanced OT problem in the discrete case. Let $\{x_i \in \mathcal{X}\}_{i=1}^n$ and $\{y_i \in \mathcal{Y}\}_{i=1}^m$. Let $\alpha = \sum_{i=1}^n \mathbf{a}_i \delta_{x_i}$ and $\beta = \sum_{i=1}^m \mathbf{b}_i \delta_{y_i}$ be discrete distributions, $\mathbf{a} \in \mathbb{R}_{>0}^n$ and $\mathbf{b} \in \mathbb{R}_{>0}^m$ with $\delta_x$ denoting the Dirac distribution on $x$. Let $\mathbf{C} \in R^{n\times m}$ be the cost matrix with $\mathbf{C}_{ij} = d(x_i, y_i)$ where $d(\cdot,\cdot) : \mathbb{R}^d \to \mathbb{R}$ denotes a distance function. The optimal coupling obtained by the regularised Optimal Transport problem reads

$$\mathbf{P}^* = \text{argmin}_{P \in \mathbf{U}(\mathbf{a},\mathbf{b})} \langle \mathbf{C}, \mathbf{P}\rangle - \epsilon H(\mathbf{P}) \tag{10}$$

where $H(\mathbf{P}) = -\sum_{i,j} \mathbf{P}_{ij}(\log(\mathbf{P}_{ij}) - 1)$ is the discrete entropy of the transport matrix. $\mathbf{U}(\mathbf{a},\mathbf{b}) = \{P \in \mathbb{R}^{n\times m} : \mathbf{a} = \mathbf{P}\mathbb{1}_m, \mathbf{P}^T\mathbb{1}_n = \mathbf{b}\}$ denotes the set of valid transport matrices.

The optimal transport matrix for the unbalanced case [Chi+18] is given by

$$\mathbf{P}^* = \text{argmin}_{P \in \mathbb{R}^{n\times m}} \langle \mathbf{C}, \mathbf{P}\rangle + \lambda_a \mathbb{D}_{KL}(\mathbf{P}\mathbb{1}_m || \mathbf{a}) + \lambda_b \mathbb{D}_{KL}(\mathbf{P}^T\mathbb{1}_n || \mathbf{b}) - \epsilon H(\mathbf{P}) \tag{11}$$

Following [Cut+22b] we define $\tau_a = \frac{\lambda_a}{\lambda_a + \epsilon}$ and $\tau_b = \frac{\lambda_b}{\lambda_b + \epsilon}$ to obtain the balanced case with $\tau_a = \tau_b = 1$. We refer to $\mathbf{P}^*\mathbb{1}_m$ as the posterior left marginals and $\mathbf{P}^{*T}\mathbb{1}_n$ as the posterior right marginals, respectively.

##### A.3 UPMM algorithm

**Algorithm 1** Unbalanced Parameterized Monge Maps

**Input** Source distribution $Q$, target distribution $P$, batch size $M$, number of inner iterations $S$, number of outer iterations $T$, regularisation parameter $\epsilon$, left unbalancedness parameter $\tau_a$, right unbalancedness parameter $\tau_b$, number of inner samples $N$

```
for t = 1, ..., T do
    Sample batch {X_i}_{i=1}^M ~ P, {Y_i}_{i=1}^M ~ Q
    Compute π_{τ_a,τ_b} = π_{τ_a,τ_b}({X_i}_{i=1}^M, {Y_i}_{i=1}^M) ∈ R_{≥0}^{M×M}
    for s = 1, ..., S do
        Draw {(k_n, l_n) ~ π_{τ_a,τ_b}}_{n=1}^N
        Compute J(θ_f, θ_g) = (1/N) Σ_{n=1}^N −f_{θ_f}(X_{k_n}) − ⟨Y_{l_n}, ∇g_{θ_g}(Y_{l_n})⟩ − f_{θ_f}(∇g_{θ_g}(Y_{l_n}))
        Update θ_g to minimize J(θ_f, θ_g)
    end for
    Update θ_f to minimize J(θ_f, θ_g)
end for
```

###### A.3.1 Model architecture

An Input Convex Neural Network (ICNN) parameterizes a function $f$ such that $f$ is convex with respect to its input by imposing certain constraints [AXK17]. Following Makkuva et al. we train two ICNNs, denoted by $f$ and $g$, with the following architecture [Mak+20]:

- $K$ *dense* layers consisting of weights $A_0, \ldots, A_K$ applied to the raw input $x$,
- $K-1$ *positive dense* layers consisting of non-negative weights $W_1, \ldots, W_K$ applied to intermediate outputs $z_{k-1}$ as defined below.

Then, layer $k$ is defined as

$$z_k = \phi((W_k z_{k-1}) + (A_k x + b_k)), \tag{12}$$

where $\phi$ is a convex non-decreasing activation function, $b_k$ the bias term and $A_k$ the weight matrix. In the last layer, we apply no activation function. Additionally, we use a quadratic first layer:

$$z_0 = (\phi(A_0 x + b_0))^2. \tag{13}$$

We enforce the non-negativity constraint on of the weights $W$ by weight clipping, while we only use a penalization term for negative weights of $g$

$$R(W^g) = \sum_{w \in W^g} ||\max(0, -w)||_2^2 \tag{14}$$

where $W^g$ denotes the set of weight matrices in the positive dense layers of the ICNN parameterizing $g$.

##### A.4 Training details

###### A.4.1 Pretraining

We pretrain the ICNN parameterizing $f$ on the identity map as suggested in [Kor+19; Amo+22] such that $\nabla f(x) = x$ and then copy the weights to $g$ for them to be mutually inverse $\nabla f(\nabla g(x)) \approx x$ and $\nabla g(\nabla f(x)) \approx x$. Therefore, we train on $X \sim \mathcal{N}(0, 3)$ for $15,000$ iterations.

###### A.4.2 Hyperparameters

For all reported experiments (both simulated data and pancreas data) we set the regularization parameter $\epsilon = 0.1$ for the computation of discrete OT in UPMM and use the following hyperparameters for training:

- learning rate: $0.001$
- optimizer: *Adam*($\beta_1 = 0.5, \beta_2 = 0.9$)

- hidden layers: $[64, 64, 64, 64]$
- inner loop iterations: $10$
- outer loop iterations: $25000$
- batch size: $1024$
- activation function: *Leaky ReLU($\beta = 0.01$)*
- gradient clipping to norm: $1.0$

Additionally, for the balanced settings, we perform best model selection based upon the lowest forward Sinkhorn divergence which is a debiased estimate of the Wasserstein distance defined as

$$SD_2^2(P,Q) = W_2^2(P,Q) - \frac{1}{2}W_2^2(P,P) - \frac{1}{2}W_2^2(Q,Q). \tag{15}$$

We also empirically observe better results using this stopping criterion for the unbalanced setting.

##### A.4.3 Influence of unbalancedness in the pancreatic endocrinogenesis data

For experiments on the pancreatic endocrinogenesis dataset, we evaluated the results obtained by different levels of unbalancedness $\tau = \tau_a = \tau_b$. For the experiments reported in Table 1 we chose $\tau = 0.85$. In Figure 3 we can see how introducing unbalancedness improves performance in correct cell type transitions.

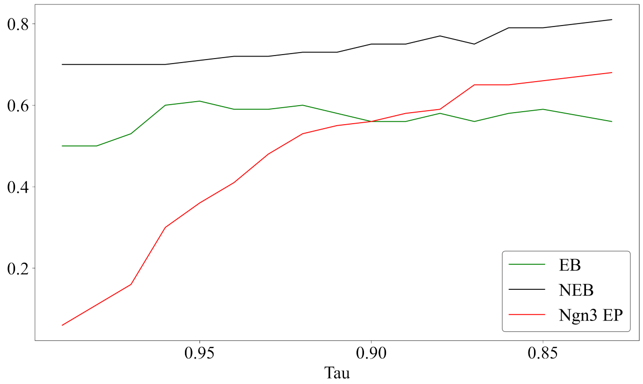


Figure 3: Effect of unbalancedness parameter $\tau$ on aggregated transition probabilities. The trend shows that introducing unbalancedness to a certain extent by lowering $\tau$ improves performance for all lineages. Metrics are computed the same way as in table 1.

Additionally, the effect of unbalancedness is visualized in Figure 4, where the changes are particularly significant for Ngn3 EP cells.

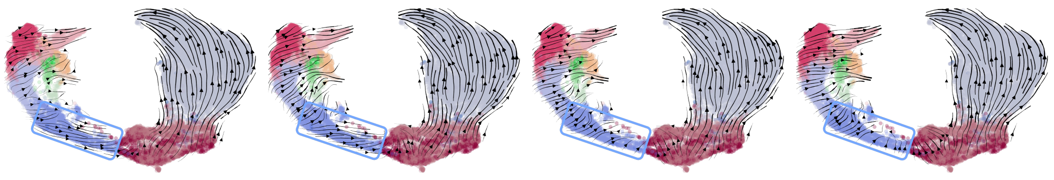

(a) PMM (b) UPMM $\tau = 0.95$ (c) UPMM $\tau = 0.90$ (d) UPMM $\tau = 0.85$

Figure 4: Effect of $\tau$ on velocity stream embedding plots.

##### A.4.4 Implementation

Our implementation is based on JAX [Bra+18], heavily building up on OTT-JAX [Cut+22a] while utilizing parts of the DeepMind JAX ecosystem [Bab+20].

##### A.5 Datasets

###### A.5.1 Simulated data

The simulated data consists of the union of draws of uniform distributions on $Q_1 \sim \mathcal{U}([-0.5, 0.5] \times [-1.5, -0.5])$ and $Q_2 \sim \mathcal{U}([4.5, 5.5] \times [-1.5, -0.5])$. Similarly, $P_1 \sim \mathcal{U}([-0.5, 0.5] \times [0.5, 1.5])$ and $P_2 = \mathcal{U}([4.5, 5.5] \times [0.5, 1.5])$. The source distribution is obtained by drawing 180 samples from $Q_1$ and 120 samples from $Q_2$. Similarly, the target distribution $P$ is obtained by 180 samples from $P_2$ and 120 samples from $P_1$.

###### A.5.2 Pancreatic endocrinogenesis data

The pancreatic endocrinogenesis data includes samples of embryonic days 14.5 and 15.5 [Bas+19]. After standard preprocessing $16,206$ genes remained. The PMM-based algorithms and Waddington OT were run on the 50 principal components. Figure 5 visualizes the distribution shift between the two time points.

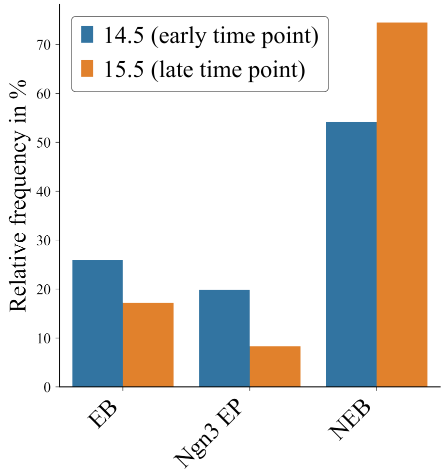


Figure 5: Distribution over the selected lineages and time points. Cells of the non-endocrine branch is much more abundant at the later time point due to very high proliferation rates of Acinar and Ductal cells.

##### A.6 Details on evaluation of competing methods

For evaluating cell type transitions we use CellRank kernels. Here, kernels quantify transition probabilities based on vector fields. For all methods yielding velocity vectors (PMM, PMM+GR, UPMM, scVelo, TrajectoryNet), we use the VelocityKernel. For evaluating Waddington OT (WOT), we use the WOTKernel.

###### A.6.1 scVelo

To infer RNA velocity with scVelo, we first selected genes measured in at least 20 cells in both unspliced and spliced transcripts. Next, cells are normalized by the median cell size, and the 2000 highly variable genes are selected. For these preprocessing steps, we used scVelo's `filter_and_normalize` function. Following, moments were calculated by the `scvelo.pp.moments` function with the settings `n_pcs=50` principal components, and `n_neighbors=30` nearest neighbors. RNA velocity was inferred using the `recover_dynamics` function implemented in scVelo.

###### A.6.2 TrajectoryNet

We run trajectory net with default parameters as suggested by the author's Jupyter notebooks. Specifically, `embedding=PCA`, `max_dim=10`, `max_iterations=10,000` and `vecint=1e-4`. We computed velocities by subtracting the inferred coordinates from the original coordinates in the embedding space. Since we were able to retrieve the inferred coordinates only for one time point, we set the velocities of the other time point to **0**.

##### A.6.3 Waddington OT

The Waddington OT results were calculated with CellRank's `WOTKernel`. For the corresponding transition matrix, we considered both inter and intra timepoint transitions. The intra timepoint transitions were quantified for each time point independently by the cell-cell nearest neighbor graph, and assigned a weight of $0.2$. Summarizing, `WOTKernel`'s `compute_transition_matrix` method was run with `growth_iters=3`, `growth_rate_key="growth_rate_init"`, `self_transitions="all"`, and `conn_weight=0.2`.

#### A.7 Velocity Stream Embedding

Velocity vectors were projected onto the two-dimensional UMAP embedding using scVelo's `velocity_embedding_stream` function. To project the high-dimensional vectors, we consider the empirical displacement given by the difference of a cellular representation in the low-dimensional embedding. The displacement vector in UMAP coordinates is then defined as the expected empirical displacement under a transition matrix and corrected by the expected shift under a uniform distribution. To define the entry $(j, k)$ of the transition matrix, consider the reference cell $j$ and a neighbor $k$. The probability that cell $j$ transitions into cell $k$ is defined as the normalized Pearson correlation between the empirical displacement in the high dimensional f of the two cells, and the velocity vector of the reference cell.

#### A.8 Cell type transition metrics

##### A.8.1 Definition

We follow [Bas+19] to obtain the ground truth of cell type transitions. We only consider cell type transitions where the target cell state is a terminal cell state $t \in T = \{$Acinar, Ductal, Alpha, Beta, Delta, Epsilon$\}$ or a union thereof. Let **ED** be the set of endocrine cell types (Alpha, Beta, Delta, Epsilon). We assume the following cell type transitions are exclusively correct (denoted by $\rightarrow$), i.e. there is no descending cell type (or set of cell types) other than the given one. We partition all considered cell type transitions into three categories.

The first set of considered transitions are endocrine branch (**ED**) transitions:

- Fev+ Alpha (**FA**) $\rightarrow$ Alpha (**A**)
- Fev+ Beta (**FB**) $\rightarrow$ Beta (**B**)
- Fev+ Delta (**FD**) $\rightarrow$ Delta (**D**)
- Fev+ Epsilon (**FE**) $\rightarrow$ Epsilon (**E**)
- **A** $\rightarrow$ **A**
- **B** $\rightarrow$ **B**
- **D** $\rightarrow$ **D**
- **E** $\rightarrow$ **E**

The second set of transitions are **Ngn3 EP** transitions:

- Ngn3 high early (**NE**) $\rightarrow$ **ED**
- Ngn3 high late (**NL**) $\rightarrow$ **ED**

The third set of transitions is the non-endocrine branch (**NEB**)

- Ductal (**DU**) $\rightarrow$ **DU**
- Tip (**T**) $\rightarrow$ Acinar (**AC**)
- **AC** $\rightarrow$ **AC**

Table 2: Cell type transition probabilities between cell types $A$ and $B$ such that cell type $A$ maps exclusively to cell type $B$. For each column, we underline the best, second best, and third best methods.

| Model | FA → A | A → A | FB → B | B → B | FD → D | D → D | FE → E | E → E | NE → ED | NL → ED | DU → DU | T → AC | AC → AC |
|---|---|---|---|---|---|---|---|---|---|---|---|---|---|
| TrajectoryNet | 0.07 | 0.46 | 0.07 | 0.39 | 0.11 | 0.75 | 0.10 | 0.52 | 0.00 | 0.01 | 0.35 | 0.79 | 0.99 |
| scVelo | **0.79** | **0.80** | **0.30** | 0.61 | 0.03 | 0.52 | 0.04 | **0.56** | **0.98** | **1.00** | 0.32 | 0.04 | 0.90 |
| WOT | **0.37** | **0.62** | 0.18 | 0.44 | 0.19 | 0.74 | **0.55** | 0.49 | 0.50 | 0.50 | **0.82** | 0.48 | 0.84 |
| PMM | **0.47** | 0.58 | 0.18 | **0.70** | **0.55** | **0.96** | **0.47** | 0.27 | 0.05 | 0.07 | 0.07 | **1.00** | **1.00** |
| PMM+GR | 0.36 | **0.73** | **0.26** | **0.72** | **0.73** | **0.98** | **0.56** | **0.68** | **0.98** | **1.00** | **0.52** | **0.94** | **1.00** |
| UPMM | 0.22 | 0.61 | **0.30** | **0.74** | **0.80** | **0.99** | 0.38 | **0.68** | **0.62** | **0.69** | **0.37** | **1.00** | **1.00** |

##### A.8.2 Detailed cell type transition results

In Table 2 we report the transition probabilities for all above-mentioned cell type transitions, the full cell type transitions to all terminal states can be found in Appendix A.13.

#### A.9 Cell type redundancy

This metric helps to understand the learned dynamics by determining macrostates that are considered to be sinks. Technically speaking, this metric looks for the minimal `n_states` − 6 in

```
cellrank.tl.terminal_states
```

such that all six terminal cell populations Acinar, Ductal, Alpha, Beta, Delta, and Epsilon are recovered. By definition, the *cell type redundancy* metric assigns a non-negative number with 0 being the best score. This metric reveals an aggregated view of the underlying cell transitions with a particular focus on less abundant cell types.

#### A.10 Velocity consistency

The *velocity consistency* metric is computed using

```
scvelo.tl.velocity_confidence
```

which measures how much the velocity of a cell correlates with the velocities of its neighbors. The neighborhood graph is given through a cell-cell neighbor graph based on transcriptomic similarity. As such, if the neighborhood is sufficiently small we expect neighboring cells to move into a similar direction. The robustness of the score has previously been reported [Ber+20].

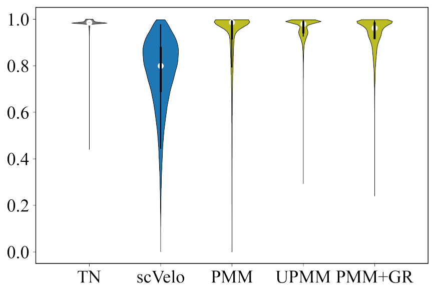


Figure 6: Velocity consistencies for all considered models which yield velocity vectors.

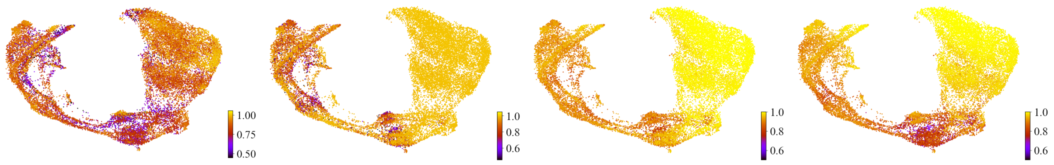


(a) scVelo (b) PMM (c) UPMM (d) PMM + GR

Figure 7: Embedding of the pancreatic endocrinogenesis data colorized by velocity consistency. While the velocity consistency of scVelo is uniformly low across all cell types, the PMM and UPMM models yield less consistent velocities in the endocrine branch. However, the consistency in the Acinar cell population is very high. We conjecture that this observation reflects the more versatile dynamics in the endocrine branch (leading to 4 different terminal cell states) while the dynamics in the Acinar cells are expected to be more uniform.

Note that there is no common way to transform the output of Waddington OT / discrete OT into velocities in gene expression space (or its embedding). Consequently, we did not apply the metric to the WOT estimates.

##### A.11 Potentials

###### A.11.1 Potentials of the simulated data

Figure 8 shows the potentials $\frac{1}{2}||y||^2 - g(y)$, defined in (4) and (5), corresponding to the transport maps plotted in figure 1. While the PMM in 8b clearly contains a sharp edge in the bottom left which defines the separation between those data points going straight to the top and those data points going across the diagonal. With increasing unbalancedness, i.e. decreasing parameter $\tau_a = \tau_b$ the angle between the vertical axis and the diagonal across which samples are mapped becomes smaller until samples are mapped vertically and the potential lines become approximately horizontal.

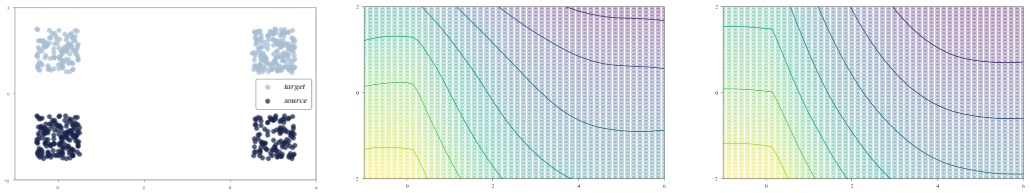


(a) Data (b) Standard Neural OT, [Mak+20] (c) $\tau = 1.0$

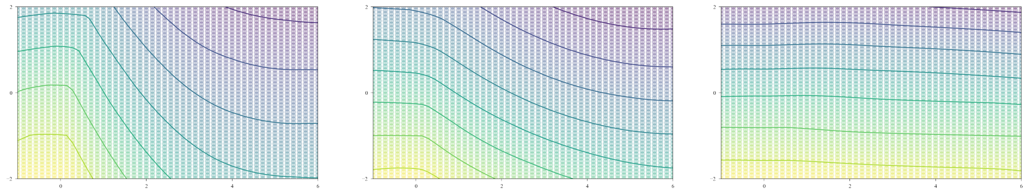

(d) $\tau = 0.99$ (e) $\tau = 0.97$ (f) $\tau = 0.9$

Figure 8: Data drawn from a mixture of uniform distribution (Appendix A.5.1). Result of PMM as proposed in [Mak+20] and UPMM with different unbalancedness parameters $\tau = \tau_a = \tau_b$

.

###### A.11.2 Potentials of the pancreatic endocrinogenesis data

Here, we evaluate the potential of each cell and plot it on a two-dimensional UMAP embedding, which allows obtaining a notion of pseudo time,

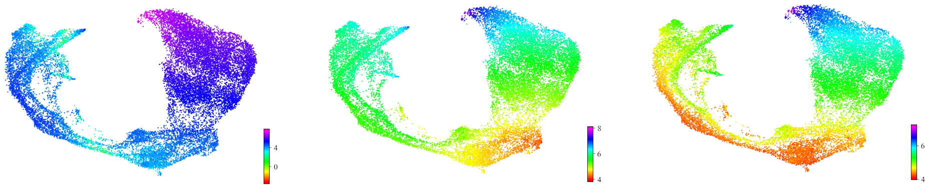

(a) PMM (b) PMM+GR (c) UPMM

Figure 9: Log-scaled potentials for PMM, PMM+GR, and UPMM on the pancreatic endocrinogenesis data (Appendix A.5.2). The darker the value, the further the cell in the developmental process.

Table 3: Evaluation of different levels of growth rate reweighting. For each column, we underline the best, second best, and third best methods.

| | Correct transitions | | |
|---|---|---|---|
| Model | **EB** | **Ngn3 EP** | **NEB** |
| PMM + GR, $c = 1$ | 0.56 | 0.48 | 0.73 |
| PMM + GR, $c = 2$ | **0.60** | **0.78** | **0.74** |
| PMM + GR, $c = 3$ | **0.59** | **0.97** | **0.79** |
| PMM + GR, $c = 4$ | **0.63** | **0.99** | **0.82** |

##### A.12 Adapted growth rates

To incorporate biological priors, we follow Waddington OT's approach of using proliferation and apoptosis marker genes. The proliferation and apoptosis scores $p_i$ and $a_i$, respectively, are obtained with Scanpy's [WAT18] `scanpy.tl.score_genes` function for each cell $x_i$. Apoptosis genes are taken from `https://www.gsea-msigdb.org/gsea/msigdb/cards/HALLMARK_P53_PATHWAY`, proliferation genes from [Tir+16].

We transform the source distribution, which by default is assumed to be uniform, such that for each cell $x_i$ the transformed mass is given by

$$P(X = x_i) = \frac{1}{Z} \exp(c\dot{(}p_i - a_i)) \tag{16}$$

with $Z$ being a normalizing constant and $c$ a parameter to be chosen. This parameter determines the magnitude of the reweighting of the masses based on the marker genes score. For the experiments reported in Table 1 we chose $c = 4$. This choice yields the best results overall. Table 3 shows the results evaluated for different $c$.

##### A.13 Full transition probabilities

In Figures 10, 11, 12, 13, 14, 15 we report the full cell type transition probabilities based on which we computed aggregated metrics. Note that some terminal cell states (on the vertical axis) are split into multiple subclusters, which is necessary to run the CellRank pipeline.

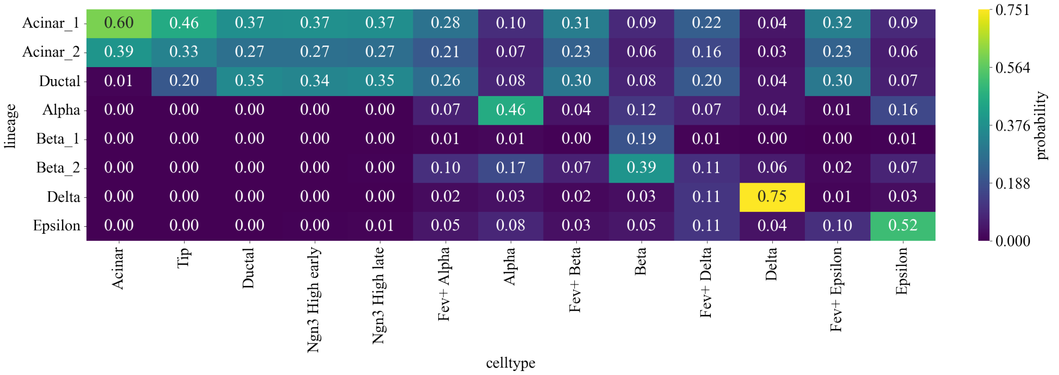


Figure 10: TrajectoryNet

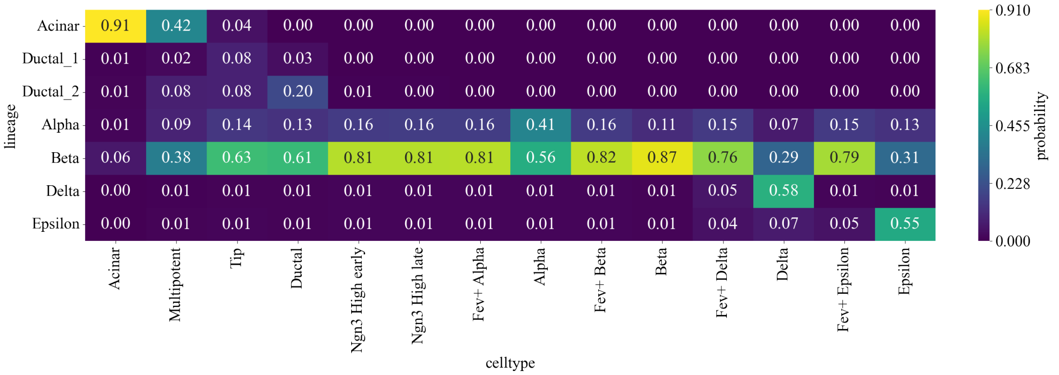


Figure 11: scVelo

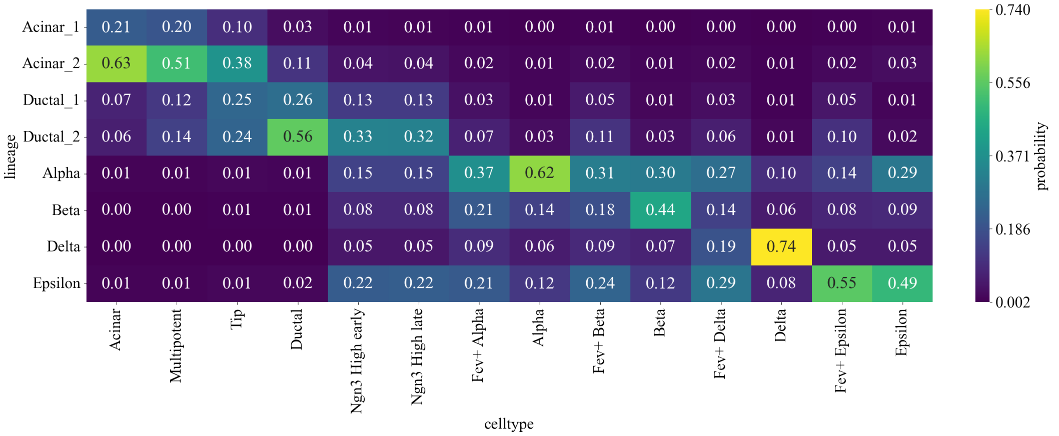


Figure 12: Waddington OT

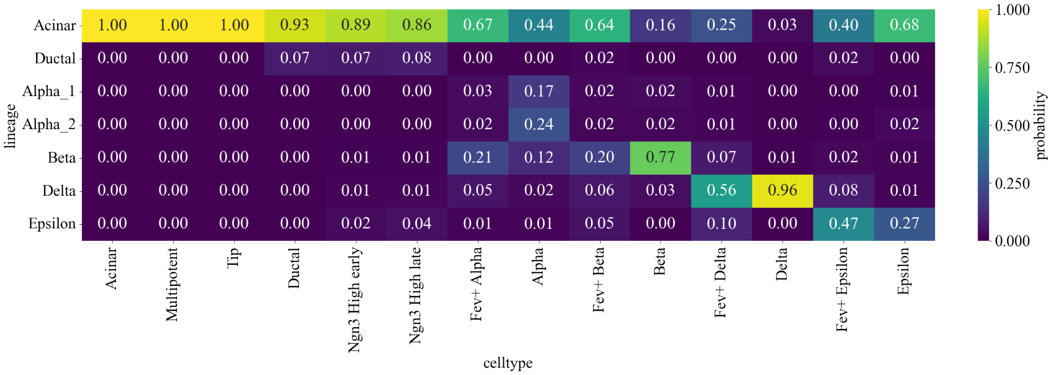


Figure 13: PMM

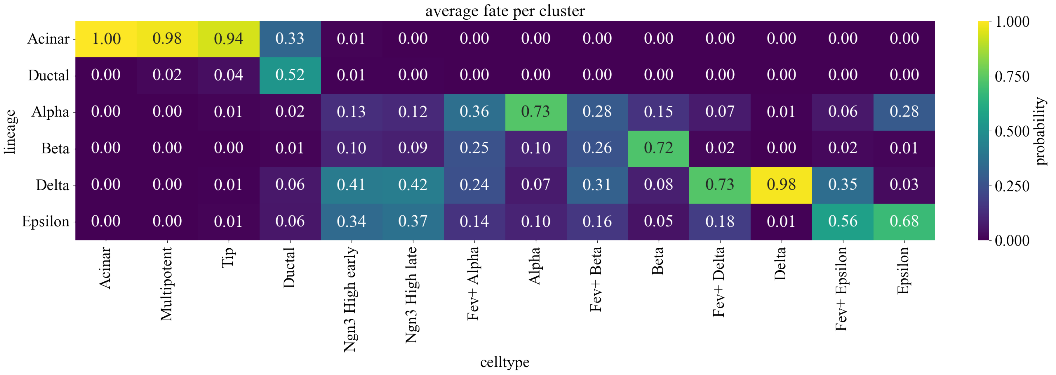


Figure 14: PMM+GR

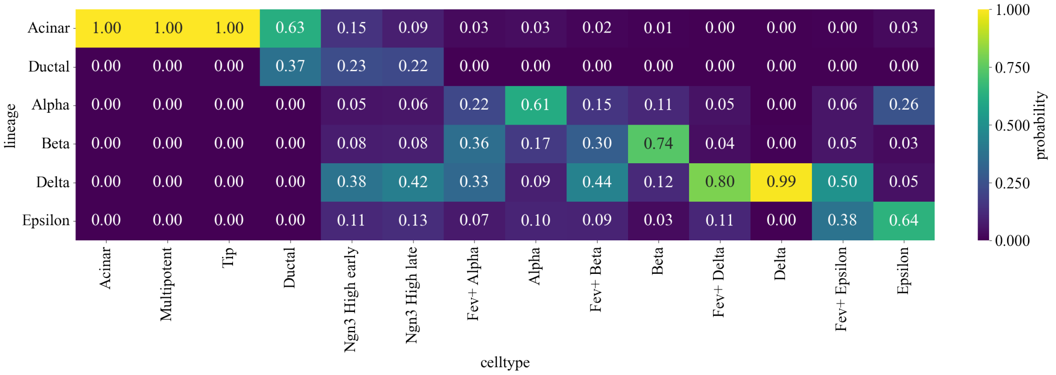


Figure 15: UPMM
